# Semiochemical responsive olfactory sensory neurons are sexually dimorphic and plastic

**DOI:** 10.1101/867044

**Authors:** Aashutosh Vihani, Xiaoyang Serene Hu, Sivaji Gundala, Sachiko Koyama, Eric Block, Hiroaki Matsunami

**Author notes:** Co-correspondence (AV), (HM). **Contributions** Conceptualization: AV and HM. Methodology: AV and HM. Investigation: AV, XSH, and HM. Formal analysis: AV. Resources: SG, SK, EB, and HM. Writing – original draft preparation: AV. Writing – review & editing: AV and HM. Funding acquisition: EB and HM. **Funding Sources:** This work was funded by NIH (DC014423 and DC016224) and NSF (1556207). Competing Interests: **None.**.

## Abstract

Understanding how genes and experiences work in concert to generate phenotypic variability will provide a better understanding of individuality. Here, we considered this in the context of the main olfactory epithelium, a chemosensory structure with over a thousand distinct cell-types, in mice. We identified a subpopulation of at least three types of olfactory sensory neurons, defined by receptor expression, whose abundances were sexually dimorphic. This subpopulation of olfactory sensory neurons was over-represented in sex-separated female mice and responded robustly to the male-specific semiochemicals 2-*sec*-butyl-4,5-dihydrothaizole and (methylthio)methanethiol. Sex-combined housing led to a robust attenuation of the female over-representation. Testing of *Bax* null mice revealed a *Bax*-dependence in generating the sexual dimorphism in sex-separated mice. Altogether, our results suggest a profound role of experience in influencing homeostatic neural lifespan mechanisms to generate a robust sexually dimorphic phenotype in the main olfactory epithelium.

## Introduction

Emergence of individuality is a ubiquitous feature across animals. It refers to the differing biological factors (“nature”), experiences (“nurture”), and randomness that generate phenotypic variability. Evidence for this variability has led to extensive research on the adaptive significance and ecological, or evolutionary, consequences of individuality^1^. Nonetheless, insight into the relative contributions and proximal mechanisms of “nature” versus “nurture” in generating phenotypic variabilities have been largely elusive.

One robust example of nature-induced inter-individual variability is sexual dimorphism. Previous work has identified both anatomical and functional substrates of this nature-induced variability in the nervous system of mice^2–5^. Similarly, extensive literature points towards an essential role of nurture-induced variability by experience-dependent, or activity-dependent, neural plasticity^6–8^. Here, to investigate how nature and nurture work in concert to generate biological variability, we focused on the mouse main olfactory epithelium (MOE), a chemosensory structure devoted to the detection of volatile odor cues. Olfactory sensory neurons (OSNs) found in the MOE express a single olfactory receptor (OR) in a monoallelic fashion out of a large and diverse family of over 1,000 candidate genes^9, 10^, thus enabling this system with an incredible potential for the emergence of individuality at the level of cell types. Odor recognition at the level of OSNs additionally has been shown to follow a combinatorial coding scheme where one OR can be activated by a set of odorants and one odorant can activate a combination of ORs^11, 12^. Through such combinatorial coding, it has been postulated that organisms, including mice and humans, can detect and discriminate against a myriad of odor molecules.

We performed RNA-Seq on the whole olfactory mucosa of mice of different sexes (“nature”) and experiences (“nurture”) to investigate origins of inter-individual differences. In doing so, we uncovered a subset of ORs that exhibit sexually dimorphic expression under sex-separated conditions. *In situ* mRNA hybridization probing for the expression of these ORs demonstrated the proportions of OSNs expressing these ORs to be over-represented in female mice. Activity-dependent labeling experiments further identified this subpopulation of OSNs as selective responders to odor cues generated by mature male mice. Targeted screening of previously identified sex-specific and sex-enriched volatiles demonstrated that this subpopulation of OSNs responded robustly to the reproductive-behavior and physiology modifying semiochemicals 2-*sec*-butyl-4,5-dihydrothaizole (SBT) and (methylthio)methanethiol (MTMT) *in vivo*. Finally, to test the role of experience in generating this sexual dimorphism, we switched male and female mice from sex-separated conditions to sex-combined conditions and learned the sexual dimorphism had severely attenuated. Examination of sex-separated mutant mice null for the BCL2-associated X protein (*Bax^-/-^)* revealed a failure to generate robust sexual dimorphisms within the whole olfactory mucosa. During the course of our investigations, a report, van der Linden et al. 2018, was also published with some overlapping findings. Altogether, these results suggest a link between specific olfactory experiences and OSN lifespan as a means to influence sensory cell-level odor representations in the olfactory system.

## Results

### Identification of sexually dimorphic ORs in the MOE

We first performed RNA-Seq on the whole olfactory mucosa of male and female mice at various ages housed under sex-separated conditions (Figure 1A). Differential expression analysis of ORs revealed no obvious sexually dimorphic OR expression at 3 weeks (weaning) age. In contrast, progressive differential expression analysis of ORs at 9, 26, and 43 weeks age revealed at least three OR genes: *Olfr910*, *Olfr912*, and *Olfr1295*, to exhibit growing and robust enrichment in female mice (Figure 1B-E). Examination of the proportion reads aligned to each of these ORs longitudinally revealed amplification of the dimorphism between the sexes with longer-term sex-separation (Figure 1F). Furthermore, this amplification appeared to exhibit receptor-specific patterns, as *Olfr1295* exhibited near-maximally dimorphic enrichment in female mice by 9 weeks age. In contrast, *Olfr910*, and *Olfr912*, which did not exhibit obvious dimorphic expression at 9 weeks age, were robustly dimorphic by 43 weeks age (at 43-weeks-old: *Olfr910* log_2_FC = 2.15, FDR < 6.89E-23; *Olfr912* log_2_FC = 2.15, FDR < 4.90E-15; *Olfr1295* log_2_FC = 2.62, FDR < 4.29E-18).

**Figure 1.**
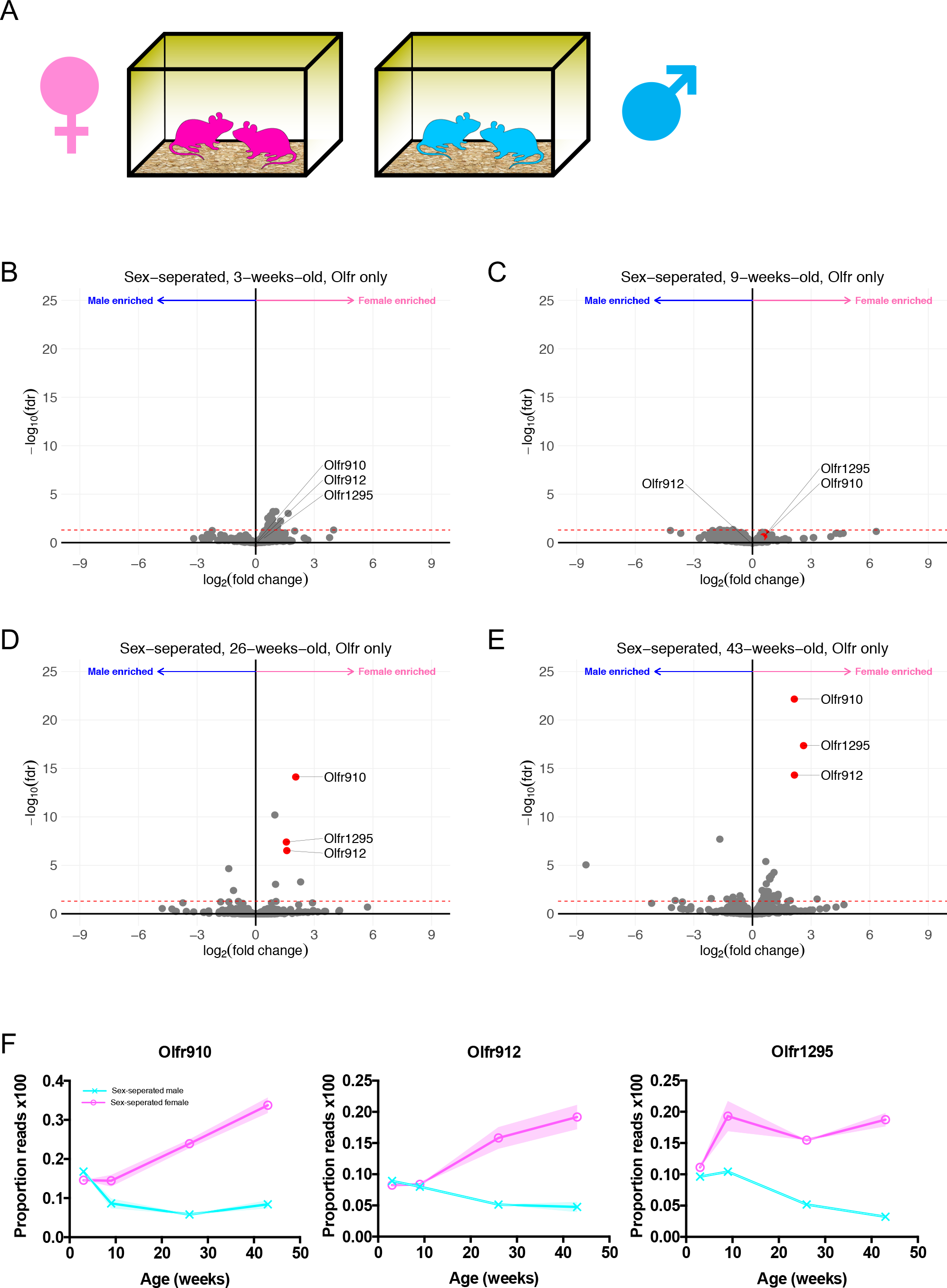
A. Schematic of the housing setup. For sex-separation, male mice were housed exclusively with male mice. Female mice were housed exclusively with female mice. B. Volcano plot comparing expression of *Olfrs* between 3-week-old sex-separated male and female mice. The red dashed line indicates an FDR = 0.05. C. Volcano plot comparing expression of *Olfrs* between 9-week-old sex-separated male and female mice. The red dashed line indicates an FDR = 0.05. D. Volcano plot comparing expression of *Olfrs* between 26-week-old sex-separated male and female mice. The red dashed line indicates an FDR = 0.05. E. Volcano plot comparing expression of *Olfrs* between 43-week-old sex-separated male and female mice. The red dashed line indicates an FDR = 0.05. F. Longitudinal plotting of the proportions of reads aligned to *Olfr910*, *Olfr912*, and *Olfr1295*. Proportions were calculated by comparing reads mapped to the specific *Olfr*s compared to those mapped to other *Olfrs*.

Past work demonstrating a correlation between OR transcript abundance and the number of OSNs expressing those ORs, led us to quantify the number of OSNs expressing these ORs by *in situ* hybridization^13, 14^. We probed specifically for the expression of *Olfr910*, *Olfr912*, and *Olfr1295* on the MOE of sex-separated male and female mice using anti-sense probes against the open reading frames (ORF) of each of these ORs. Given the 96% nucleotide sequence identity of *Olfr910* and *Olfr912*, anti-sense ORF probes against either *Olfr910* or *Olfr912* labelled OSN populations expressing either receptor (hereafter denoted as Olfr910/912). The results of the *in situ* hybridization demonstrated that the proportion of OSNs expressing *Olfr910/912* (p < 0.0001, unpaired two-tailed t-test), and *Olfr1295* (p < 0.01, unpaired two tailed t-test), was greater in 43-week-old female mice than in 43-week-old male mice (Figure 2A-D). Altogether, these results lead us to conclude that the subpopulation of OSNs expressing *Olfr910/912* and *Olfr1295* are over-represented in sex-separated female mice compared to sex-separated male mice.

**Figure 2.**
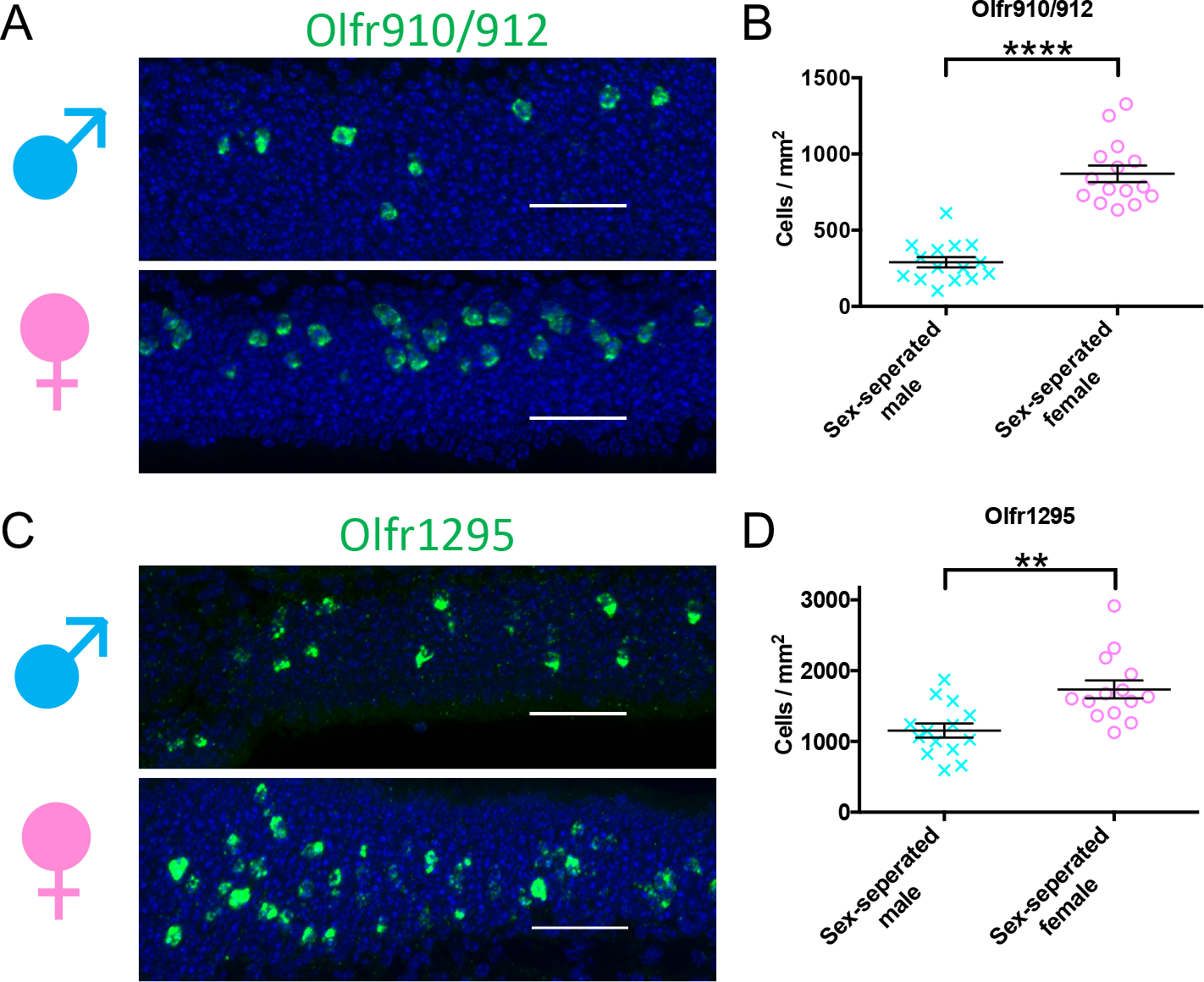
A. Representative *in situ* hybridization picture probing for the expression of *Olfr910/912* in 43-week-old sex-separated male (top) and female (bottom) mice. Scale bars indicate 50 μm. B. Summary data showing the proportion of OSNs expressing *Olfr910/912* between 43-week-old male and female mice. An unpaired two-tailed t-test revealed statistical difference (p < 0.0001) between males and females. C. Representative *in situ* hybridization picture probing for the expression of *Olfr1295* in 43-week-old sex-separated male (top) and female (bottom) mice. Scale bars indicate 50 μm. D. Summary data showing the proportion of OSNs expressing *Olfr1295* between 43-week-old male and female mice. An unpaired two-tailed t-test revealed statistical difference (p < 0.01) between males and females.

### Sexually dimorphic OSNs are selectively activated by the scent of adult male mice *in vivo*

Our identification of a subpopulation of OSNs to be over-represented in sex-separated female mice led us to hypothesize an essential role of the associated ORs in detecting sex-specific odors. To test this hypothesis, we briefly exposed juvenile mice (∼3-weeks-old) to either a blank odor cassette (control), mature male mice, mature female mice, or 10μL of 1% (v/v) acetophenone spotted onto blotting paper placed inside an odor cassette (Figure 3A). We specifically focused on OSNs expressing *Olfr910/912* or *Olfr1295* by *in situ* hybridization and performed immunostaining for the phosphorylation of ribosomal subunit S6 (pS6), a pan-neuronal marker of activity^11, 15^.

**Figure 3.**
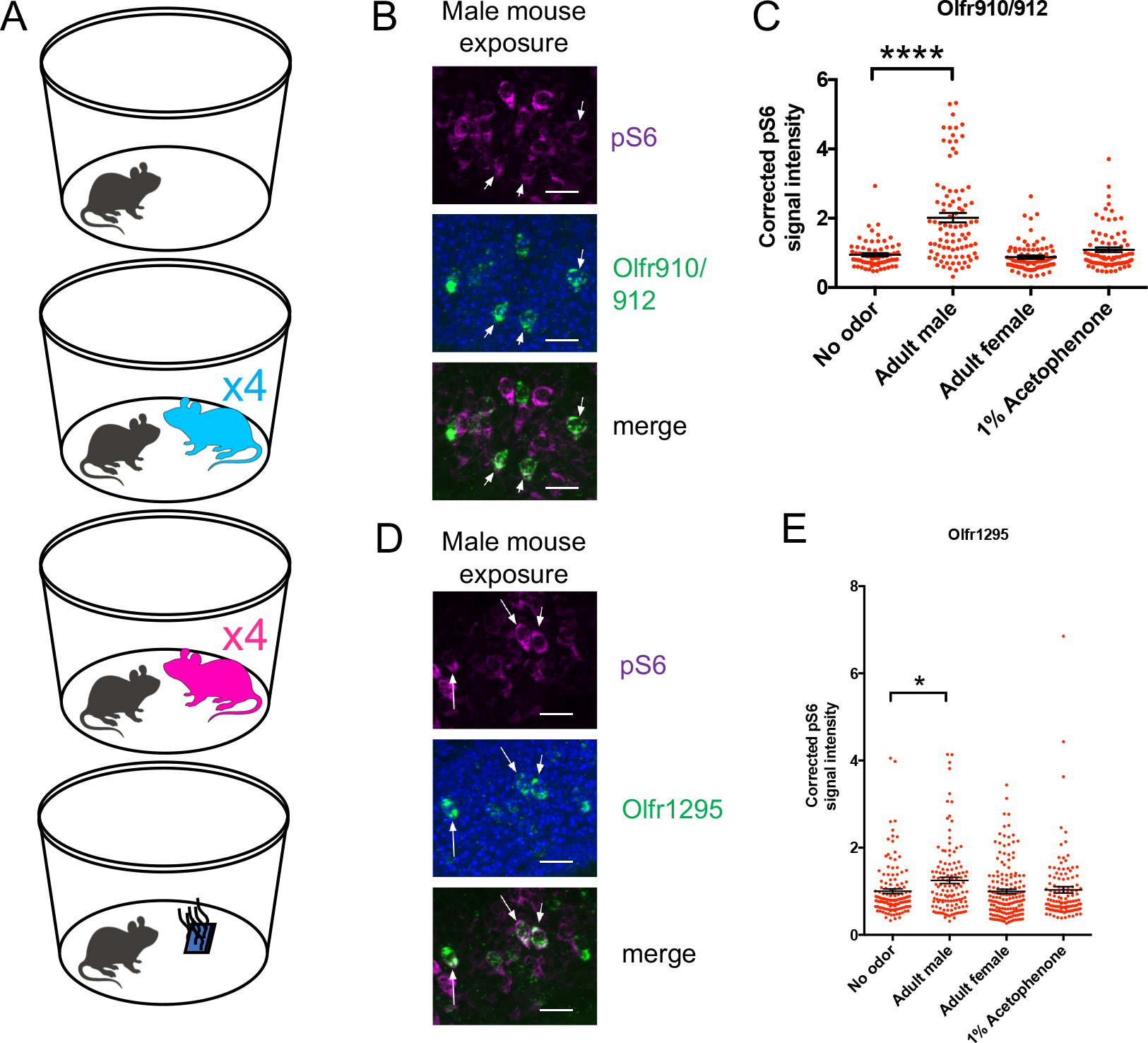
A. Schematic of exposure experiment. A juvenile mouse (black) was exposed to (in descending order) a clean environment, four adult male mice, four adult female mice, or 1% (v/v) acetophenone for 1 hour in a sealed container. B. Representative *in situ* hybridizations and pS6 immunostaining showing colocalization events between cells expressing *Olfr910/912* and pS6 signal induction following exposure of a juvenile mouse to adult male mice. Scale bars indicate 20 μm. C. Summary data showing pS6 induction in cells expressing *Olfr910/912* following exposure of a juvenile mouse to multiple stimuli. One-way ANOVA with Dunnett’s multiple comparisons test correction reveals only exposure to the adult male to lead to significant (p < 0.0001) pS6 induction within cells expressing *Olfr910/912*. D. Representative *in situ* hybridizations and pS6 immunostaining showing colocalization events between cells expressing *Olfr1295* and pS6 signal induction following exposure of a juvenile mouse to adult male mice. Scale bars indicate 20 μm. E. Summary data showing pS6 induction in cells expressing *Olfr1295* following exposure of a juvenile mouse to multiple stimuli. One-way ANOVA with Dunnett’s multiple comparisons test correction reveals only exposure to the adult male to lead to significant (p < 0.05) pS6 induction within cells expressing *Olfr1295*.

Double *in situ* hybridization and immunostaining revealed the subpopulation of OSNs expressing *Olfr910/912* (p < 0.0001, one-way ANOVA with Dunnett’s multiple comparisons test correction) and *Olfr1295* (p < 0.05, one-way ANOVA with Dunnett’s multiple comparisons test correction) to display elevated pS6 staining intensity when exposed to mature male mice (Figure 3B-E). Exposure to mature female mice or acetophenone did not lead to significant induction of pS6 in the subpopulation of OSNs expressing either *Olfr910/912* or *Olfr1295*. Altogether, our results suggest that the subpopulation of OSNs that express these receptors be selective responders to the natural scent of mature male mice.

### Sexually dimorphic OSNs are selectively responsive to specific semiochemicals *in vivo*

The observation that OSNs expressing *Olfr910/912* and *Olfr1295* are activated by the scent of mature male mice led us to hypothesize that this subpopulation of OSNs responds specifically to sexually dimorphic odors produced by mature male mice. To test this hypothesis, we searched the literature for known sex-specific or sex-enriched odors and leveraged phosphorylated S6 ribosomal subunit capture (pS6-IP) as a means to determine the molecular identities of the OSNs activated by monomolecular odorants *in vivo*. Immunoprecipitation of phosphorylated ribosomes from activated neurons followed by associated mRNA profiling by RNA-Seq (pS6-IP-Seq) and differential expression analysis, enabled us to perform an unbiased identification of the molecular profiles, including ORs expressed, of OSNs activated by specific odorants (Figure 4A)^11^.

**Figure 4.**
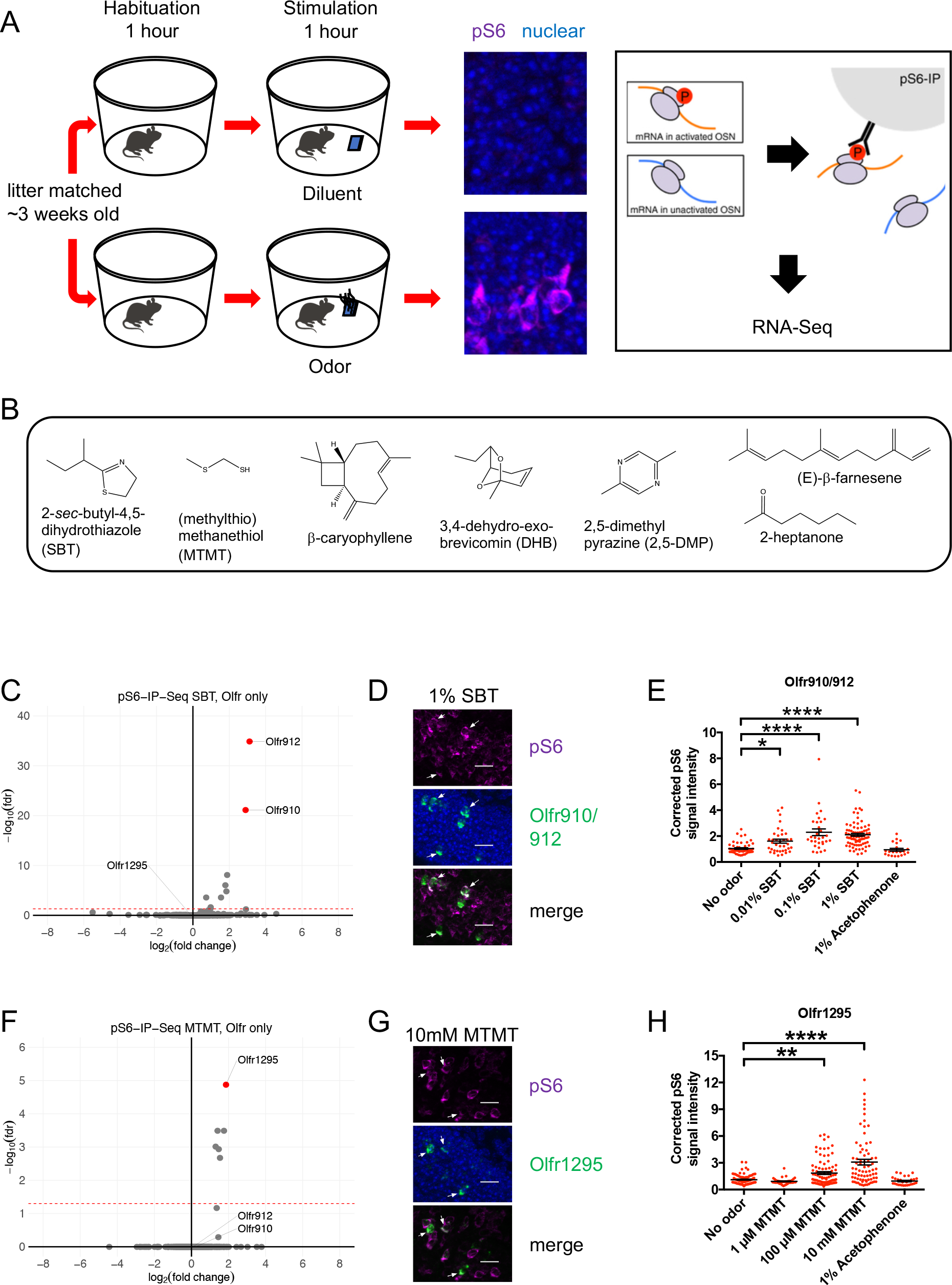
A. Schematic of the pS6-IP-Seq experiment. Litter matched, ∼3-week-old mice are used. Mice are habituated to an odor-free environment for one hour. One mouse then receives exposure to an odor stimulus, while another receives exposure to the diluent for another hour. Olfactory tissues are then harvested and immunoprecipitated using an antibody against pS6. B. The battery of sex-specific and sex-enriched volatiles screened using pS6-IP-Seq. C. Volcano plot showing the results of pS6-IP-Seq using 0.01% (v/v) SBT diluted in water as stimulus. The red dashed line indicates an FDR = 0.05. D. Representative *in situ* hybridizations and pS6 immunostaining showing colocalization events between cells expressing *Olfr910/912* and pS6 signal induction following exposure of a juvenile mouse to 1% (v/v) SBT diluted in water. Scale bars indicate 20 μm. E. Summary data showing pS6 induction in cells expressing *Olfr910/912* following exposure of a juvenile mouse to increasing concentrations of SBT and acetophenone. One-way ANOVA with Dunnett’s multiple comparisons test correction reveals only exposure to 0.01% SBT, 0.1% SBT, and 1% SBT to lead to significant pS6 induction within cells expressing *Olfr910/912* (**** p < 0.0001, * p < 0.05). F. Volcano plot showing the results of pS6-IP-Seq using 100 μM MTMT dissolved in ethanol as stimulus. The red dashed line indicates an FDR = 0.05. G. Representative *in situ* hybridizations and pS6 immunostaining showing colocalization events between cells expressing *Olfr1295* and pS6 signal induction following exposure of a juvenile mouse to 10mM MTMT diluted in ethanol. Scale bars indicate 20 μm. H. Summary data showing pS6 induction in cells expressing *Olfr1295* following exposure of a juvenile mouse to increasing concentrations of MTMT and acetophenone. One-way ANOVA with Dunnett’s multiple comparisons test correction reveals only exposure to 100μM MTMT, and 10mM MTMT to lead to significant pS6 induction within cells expressing *Olfr1295* (**** p < 0.0001, ** p < 0.01).

While the origins of the differences between the scents of mature male and female mice are diverse, we reasoned mouse urine to be a significant source of odor cues. Past literature contrasting intact male mouse urine, castrated male mouse urine, and female mouse urine volatiles has identified many components to differ in their presence and abundance^16–21^. Using pS6-IP-Seq we systematically screened the mouse urine constituents: 2-*sec-*butyl-4,5-dihydrothiazole (SBT); (methylthio)methanethiol (MTMT); β-caryophyllene; 3,4-dehydro-*exo*-brevicomin; 2,5-dimethylpyrazine; (E)-β-farnesene; and 2-heptanone (Figure 4B).

Differential expression analysis following pS6-IP-Seq across the tested panel of odorants (Figure 4C, Supplementary Figure 1, Supplementary Figure 2) led to the identification of SBT as an activator for OSNs expressing *Olfr910* and *Olfr912* and MTMT as an activator for OSNs expressing *Olfr1295*. Indeed, exposure to 10μL of 0.01% (v/v) SBT lead to the lowest false discovery rate (FDR) and greatest enrichment of transcripts for *Olfr910* and *Olfr912* from activated OSNs (at an FDR < 0.05), compared to all other ORs, suggesting OSNs expressing *Olfr910* and *Olfr912* to be the most robustly responding OSNs for SBT *in vivo* (Figure 4C) (*Olfr910* log_2_FC = 3.11, FDR < 1.25E-35; *Olfr912* log_2_FC = 2.89, FDR < 7.97E-22). Similarly, exposure to 10μL of 100μM MTMT lead to the lowest FDR and greatest enrichment of transcripts for *Olfr1295* from activated OSNs (at an FDR < 0.05), compared to all other ORs, suggesting OSNs expressing *Olfr1295* to be the most robustly responding OSNs for MTMT *in vivo* (Figure 4F) (log_2_FC = 1.86, FDR < 1.34E-5).

To independently validate the pS6-IP-Seq data, we briefly exposed juvenile mice to either an empty odor cassette (control), varying concentrations of SBT or MTMT, or 1% (v/v) acetophenone, and then harvested MOE for staining. *In situ* hybridization probing for the expression of *Olfr910/912* and immunostaining for pS6 showed OSNs expressing *Olfr910/912* to display increasingly intense pS6 immunostaining following exposure to SBT in a concentration-dependent manner (0.01% SBT p < 0.05; 0.1% SBT p < 0.0001; 1% SBT p < 0.0001, one-way ANOVA with Dunnett’s multiple comparisons test correction). Further, in accordance with our previous findings, exposure to acetophenone did not lead to significant pS6 induction in OSNs expressing *Olfr910/912* (Figure 4D-E)^11, 22^. Similarly, OSNs expressing *Olfr1295*, identified by *in situ* hybridization, displayed increasingly intense pS6 immunostaining following exposure to MTMT in a concentration-dependent manner (100 μM MTMT p < 0.01; 10 mM MTMT p < 0.0001, one-way ANOVA with Dunnett’s multiple comparisons test correction). OSNs expressing *Olfr1295* displayed a non-significant pS6 signal intensity difference following exposure to acetophenone compared to control conditions (Figure 4G-H).

To further validate our identified ligand-receptor pairs we tested mature male and female mice. We provided a brief exposure to sex-separated, 26-week-old adult male and female mice to either an empty odor cassette (control), 0.1% (v/v) SBT, varying concentrations of MTMT, or 1% (v/v) acetophenone, and then harvested MOE for staining. Double *in situ* hybridization and pS6 immunostaining revealed the following: OSNs expressing *Olfr910/912* showed pS6 signals only following SBT exposure (Figure 5A) (p < 0.0001, one-way ANOVA with Tukey’s multiple comparisons test correction), OSNs expressing *Olfr983* showed pS6 signals only following acetophenone exposure (Figure 5C) (p < 0.0001, one-way ANOVA with Tukey’s multiple comparisons test correction)^11^, and OSNs expressing *Olfr1295* showed pS6 signals only following MTMT exposure (Figure 5B) (p < 0.0001, one-way ANOVA with Tukey’s multiple comparisons test correction). We did not observe any sex bias in the responsivity of sensory cell populations at the single-cell level by pS6 signal intensity induction by exposure to either SBT or acetophenone. Female mouse OSNs expressing *Olfr1295* exhibited a subtle but significant response to 10mM MTMT compared to male mouse OSNs expressing *Olfr1295* worthy of potential future investigation (p < 0.05, one-way ANOVA with Tukey’s multiple comparisons test correction). Nonetheless, our combination of pS6-IP-Seq and *in situ* results are consistent with the idea that SBT activates OSNs expressing *Olfr910/912* more robustly than any other OSN *in vivo*, and MTMT activates OSNs expressing *Olfr1295* more robustly than any other OSN *in vivo*.

**Figure 5.**
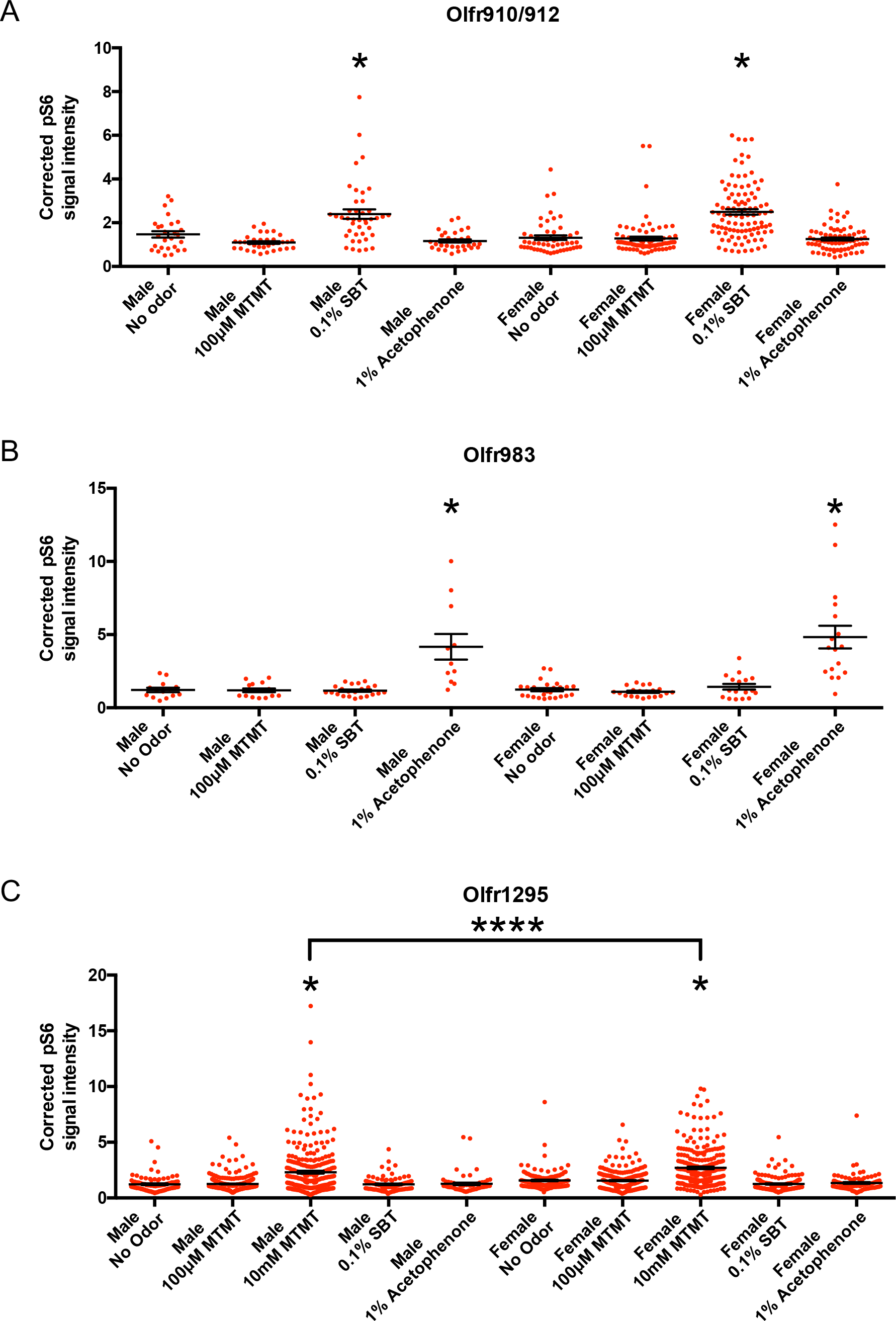
A. Comparison of responses of 26-week-old male and female mouse OSNs expressing *Olfr910*/*912* to various stimuli. One-way ANOVA with Tukey’s multiple comparisons test correction reveals only exposure to 0.1% SBT lead to significant pS6 induction (p < 0.0001) with no significant differences in male versus female responses. B. Comparison of responses of 26-week-old male and female mouse OSNs expressing *Olfr983* to various stimuli. One-way ANOVA with Tukey’s multiple comparisons test correction reveals only exposure to 1% acetophenone lead to significant pS6 induction (p < 0.0001) with no significant differences in male versus female responses. C. Comparison of responses of 26-week-old male and female mouse OSNs expressing *Olfr1295* to various stimuli. One-way ANOVA with Tukey’s multiple comparisons test correction reveals only exposure to 10mM MTMT lead to significant pS6 induction (p < 0.0001). A subtle but significantly greater response was observed in the female compared to the male (p < 0.05).

### Long-term cohabitation with the opposite sex is sufficient to attenuate sexual dimorphism in the MOE

Emerging literature has evidenced a role for experience in influencing sensory-cell representations within the olfactory epithelium^14, 23, 24^. Thus, our identification of a subset of over-represented ORs in sex-separated female mice led to us to hypothesize a role for experience in generating this dimorphism.

We hypothesized that regular exposure of a female mouse to the semiochemicals SBT and MTMT, by cohabitation with a male mouse, would influence the population dynamics of OSNs expressing *Olfr910*, *Olfr912*, and *Olfr1295*. To test this hypothesis, we used mice that were first sex-separated from weaning (∼3 weeks age) until 26 weeks of age. These sex-separated mice were then switched to cohabitation with the opposite sex (sex-combined housing) from 26 weeks age to 43 weeks age (Figure 6A). At 43 weeks age, whole olfactory mucosa from the male and female mice was harvested and processed for sequencing and histology.

**Figure 6.**
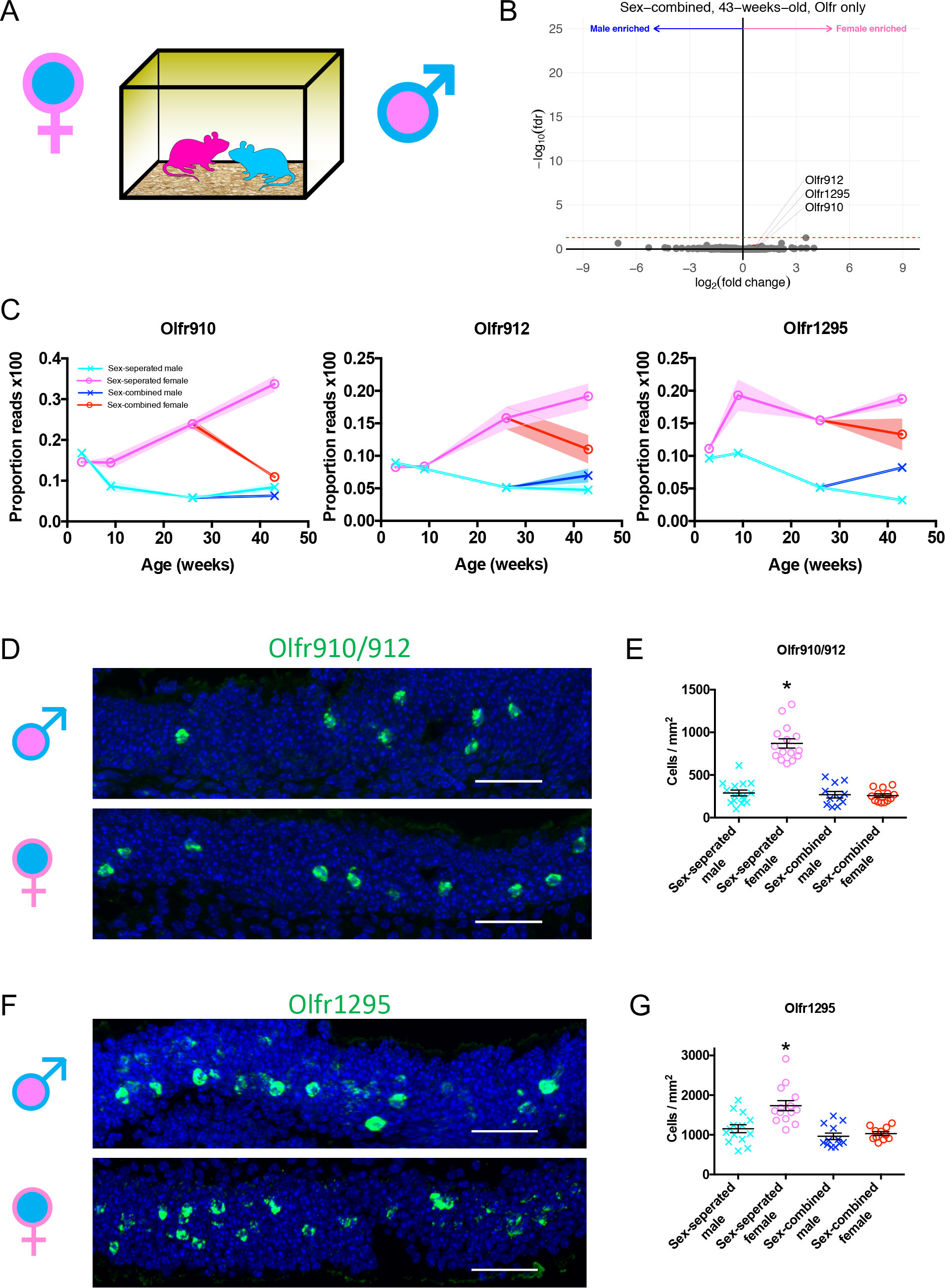
A. Schematic of the housing setup. For sex-combined housing, one male mouse was housed with one female mouse. B. Volcano plot comparing expression of *Olfrs* between 43-week-old sex-combined male and female mice. The red dashed line indicates an FDR = 0.05. C. Longitudinal plotting of the proportions of reads aligned to *Olfr910*, *Olfr912*, and *Olfr1295* of male and female mice comparing sex-separation and sex-combined housing. D. Representative *in situ* hybridization picture probing for the expression of *Olfr910/912* in 43-week-old sex-combined male (top) and female (bottom) mice. Scale bars indicate 50 μm. E. Summary data showing the proportion of OSNs expressing *Olfr910/912* between 43-week-old male and female mice housed either in a sex-separated or sex-combined fashion. One-way ANOVA with Tukey’s multiple comparisons test correction reveals only sex-separated female mice to differ in the proportions of OSNs expressing *Olfr910/912* from the others (p < 0.0001, all comparisons). F. Representative *in situ* hybridization picture probing for the expression of *Olfr1295* in 43-week-old sex-combined male (top) and female (bottom) mice. Scale bars indicate 50 μm. G. Summary data showing the proportion of OSNs expressing *Olfr1295* between 43-week-old male and female mice housed either in a sex-separated or sex-combined fashion. One-way ANOVA with Tukey’s multiple comparisons test correction reveals only sex-separated female mice to differ in the proportions of OSNs expressing *Olfr910/912* from the others (p < 0.001, sex-separated male vs sex-separated female, p < 0.0001 all others).

Differential expression analysis of whole olfactory mucosa from sex-combined male and female mice revealed a severe attenuation of the dimorphic expression of *Olfr910*, *Olfr912*, and *Olfr1295* (Figure 6B) (at 43-weeks-old: *Olfr910* log_2_FC = 0.77, FDR > 0.80; *Olfr912* log_2_FC = 0.58, FDR > 0.83; *Olfr1295* log_2_FC = 0.74, FDR > 0.82). After 17 weeks of sex-combined housing, the proportional expression of each of these receptors changed much more profoundly in female mice than male mice, again, in a receptor-specific fashion. Normalized expression of *Olfr910* and *Olfr912* appeared to be more similar between sex-separated males, sex-combined males, and sex-combined females while being distinct and less than sex-separated females. On the other hand, the normalized expression of *Olfr1295* appeared to be greatest in sex-separated females, decreasing in sex-combined females, sex-combined males, and lowest in sex-separated males (Figure 6C). *In situ* hybridization to assess the proportional abundance of OSNs expressing these receptors revealed, again, that sex-separated female mice were distinct from sex-combined female, sex-combined male, and sex-separated male mice. Sex-separated female mice had an over-representation of OSNs expressing *Olfr910/912* (p < 0.0001, one-way ANOVA with Tukey’s multiple comparisons test correction) and *Olfr1295* (p < 0.001, one-way ANOVA with Tukey’s multiple comparisons test correction) while mice from other conditions were all comparable. These findings suggest cohabitation with the opposite sex, and potentially olfactory experience, is sufficient to attenuate the over-representation of the subpopulation of sexually dimorphic OSNs (Figure 6D-G, Supplementary Figure 3).

### Generating robust sexual dimorphisms in the MOE is *Bax*-dependent

Our observation that specific experiences influence OSN population dynamics led to us hypothesize a link between OSN activity and neuronal lifespan^25, 26^. Significant past literature has evidenced a role for neural activity to lengthen the lifespan of a neuron in a Bcl2-associated X protein (Bax)-dependent manner^27^.

To test the hypothesis of OSN activity influencing OSN lifespan in a *Bax*-dependent manner, we used *Bax^-/-^* mice^27^. Compared to wild-type mice, differential expression analysis of RNA sequenced whole olfactory mucosa tissues from sex-separated 26-week-old *Bax^-/-^* male and female mice revealed an overall lack of the sexually dimorphic expression of *Olfr910*, *Olfr912*, and *Olfr1295* (Figure 7A). Additionally, *in situ* hybridization probing for the proportional abundance of OSNs expressing *Olfr910/912* and *Olfr1295* demonstrated non-significant differences in sex-separated *Bax^-/-^* male and female mice by 43 weeks age (Figure 7B-E). Altogether, these results suggest that OSN activity, by olfactory experience, influences OSN population dynamics to ultimately sculpt and shape OR representations by altering sensory neuron lifespans.

**Figure 7.**
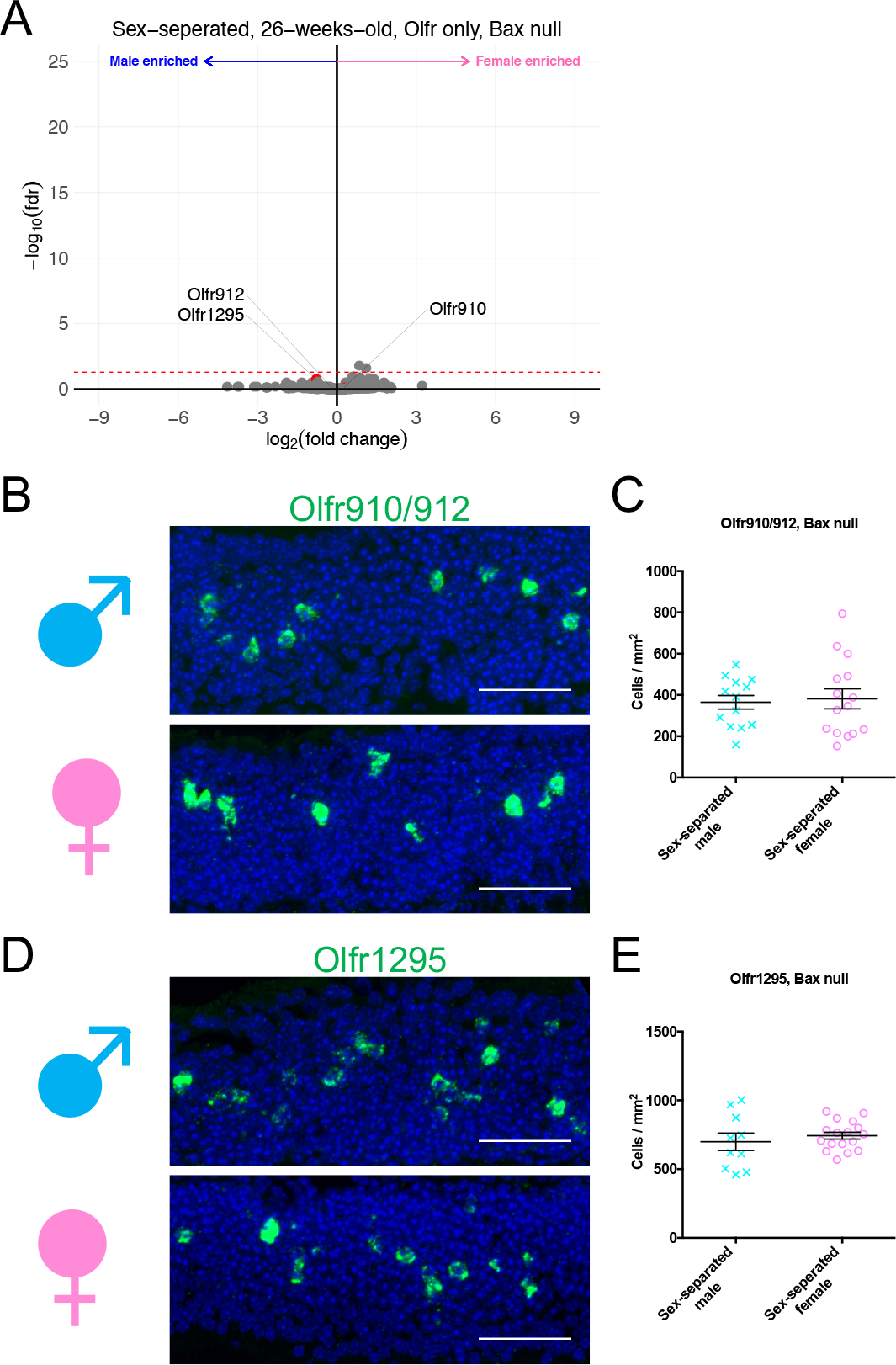
A. Volcano plot comparing expression of *Olfrs* between 26-week-old, *Bax* null, sex-separated male and female mice. The red dashed line indicates an FDR = 0.05. B. Representative *in situ* hybridization picture probing for the expression of *Olfr910/912* in 43-week-old, *Bax* null, sex-separated male (top) and female (bottom) mice. Scale bars indicate 50 A. μm. C. Summary data showing the proportion of OSNs expressing *Olfr910/912* between 43-week-old, *Bax* null, sex-separated male and female mice. An unpaired two-tailed t-test reveals no statistical difference (p > 0.05) between males and females. D. Representative *in situ* hybridization picture probing for the expression of *Olfr1295* in 43-week-old, *Bax* null, sex-separated male (top) and female (bottom) mice. Scale bars indicate 50 μm. E. Summary data showing the proportion of OSNs expressing *Olfr1295* between 43-week-old, *Bax* null, sex-separated male and female mice. An unpaired two-tailed t-test reveals no statistical difference (p > 0.05) between males and females.

## Discussion

Using a series of complimentary but independent approaches, we have identified a subpopulation of OSNs, defined by specific OR expression, to exhibit sexual dimorphism and experience-dependent plasticity. We have identified female mice, in the absence of a male, exhibit an over-representation of OSNs expressing *Olfr910/912*, and *Olfr1295*. Long-term cohabitation of a female mouse with a male mouse led to the attenuation of these over-represented ORs. To confirm an OSN activity-dependent component of this experience, we demonstrate this subpopulation of OSNs is not only activated by the natural scent of mature male mice, but is also exquisitely and robustly responsive to the previously identified male-specific semiochemicals SBT and MTMT. Finally, the observation that sex-separated *Bax^-/-^* mice fail to generate robust sexual dimorphisms suggest a role of cell death in generating our observed differences in the proportional number of OSNs expressing *Olfr910/912* and *Olfr1295* in sex-separated male and female mice.

### Olfactory experience as a mechanism to influence OSN population dynamics

Given the capacity of the MOE to regenerate throughout the life of an animal^28^, it has been suggested activity-mediated mechanisms may “individualize” the olfactory system by influencing OSN population dynamics^14^. While we acknowledge cohabitation of the opposite sexes induces a number of changes in the nervous system of mice (compared to sex-separation)^2, 3^, we propose that changes in OSN population dynamics, following cohabitation, for OSNs expressing *Olfr910/912* and *Olfr1295*, are in part mediated by olfactory experience with SBT and MTMT. The lengthy timeframes necessary to generate the differences that we observe are consistent with the hypothesis of modulation of OSN lifespan by activity. Nonetheless, we cannot rule out contributions of experience on OSN neurogenesis rates or OR gene choice. Questions regarding the individual contributions, or non-contributions, of OSN development and lifespan to generate this dimorphism remain open.

Additionally, the form of plasticity we observe appears to employ a distinct mechanism from reports of fear conditioning influencing the proportional abundance of OSNs expressing ORs responsive to the fear-conditioned odor^29–31^. In the case of fear conditioning to acetophenone, the number of OSNs expressing M71 appears to profoundly upregulate within just 3 weeks. In contrast, even by 9 weeks (∼6 weeks post-weaning), the difference in the expression of *Olfr910* and *Olfr912* appear to be insignificant in male versus female mice. Altogether, these observations lead us to speculate the existence of a multitude of distinct mechanisms, operating at non-identical time scales, to influence OSN population dynamics. Future work to identify and demonstrate these mechanisms is necessary to deepen our understandings of these phenomena and experience-dependent plasticity.

### A decrease in male-specific semiochemical responsive OSNs in females following sex-combined housing

The finding of over-represented OSN subpopulations robustly responsive to male-specific semiochemicals, in sex-separated females, to decrease in proportional abundance following sex-combined housing, while consistent with recent literature^14, 23, 24^, is unexpected. The results suggest that once females receive exposure to male-specific semiochemicals, their detectability for semiochemicals slowly decreases over time, reflecting a potential homeostatic “gain control” mechanism for salient cue detection at the level of primary sensory neurons.

Our further finding that sex-separated *Bax^-/-^* mice to not exhibit sexual dimorphism point towards a potential role of activity-dependent changes in neural lifespan in generating sexually dimorphic OR expression in sex-separated wild-type mice. Though *Bax* would certainly exert experience-independent effects in a mouse, we speculate the lack of robust sexual dimorphism in sex-separated Bax^-/-^ mice to be a result of the lack of *Bax*-regulated activity-dependent alteration of sensory neuron lifespan.

### Sexual dimorphism in OSNs responsive to semiochemicals

Plasticity within the OSN population can be postulated to enable adaptation of an individual’s olfactory system for the sensitive detection of salient odors, which may vary from one environment to another. While sex-specific chemical cues have been implicated in instinctual behaviors and physiology^16, 20, 32–38^, the degree to which animals are exposed to these chemical cues in nature may vary substantially among individuals.

A recent report by van der Linden et al. 2018 also identified sexually dimorphic expression of a subset of ORs using a combination of sequencing and histology-based approaches^24^. Our data agree in the following manner: identification of the sexually dimorphic expression of *Olfr910*, *Olfr912*, *Olfr1295*, *and Olfr1437* in mice housed in a sex-separated manner, demonstration of activation of OSNs expressing *Olfr910, Olfr912,* and *Olfr1295* following exposure to mature male mice, and a general lack of sexually dimorphic ORs in mice housed in a sex-combined manner. Together, our findings of experience to influence OSN population dynamics suggest a role in adjusting an animal’s sensitivity to the salient chemical cues of male mice.

Our identification of the semiochemicals SBT and MTMT as robust agonists for *Olfr910*, *Olfr912*, and *Olfr1295* posit a number of intriguing speculations. Remarkably, other ORs activated by SBT and MTMT did not exhibit sexual dimorphism. Furthermore, when we tested other sex-specific and sex-enriched odorants, we did not observe activation of OSNs expressing *Olfr910*, *Olfr912*, or *Olfr1295* (Supplementary Figure 1), nor did cognate receptors for these other odors exhibit sexually dimorphic expression (Supplementary Figure 2). These results altogether lead us to hypothesize a specialized role for *Olfr910*, *Olfr912*, and *Olfr1295* in conveying a salient signal from the olfactory periphery to the central nervous system.

The identification of a subpopulation of OSNs to also be plastic and robustly responsive to male-specific semiochemicals also raise speculations about the flexibility of an individual’s behavioral responses to semiochemicals. That is, while behavioral and physiological responses to semiochemicals have traditionally been viewed as genetically predetermined and “hardwired”, there may exist a significant context and experience-dependent flexibility. For example, it has been previously shown that group housed sex-separated female mice exhibit a general suppression and irregularity in estrous cycling. Upon exposure to a mature male mouse, these unisexually grouped female mice often rapidly and synchronously enter into estrus (Whitten effect)^39–43^. Past implications of SBT also inducing the Whitten effect^16^, and our finding of *Olfr910* and *Olfr912* to be robustly responsive to SBT and over-represented in sex-separated female mice, lead us to speculate that the over-representation of these ORs to serve to enhance SBT detection for mediation of the Whitten effect. Past implications of MTMT influencing female mouse attractive behaviors^18^, and our finding of *Olfr1295* to be robustly responsive to MTMT and over-represented in sex-separated female mice, lead us to speculate over-representation of this OR to serve to enhance MTMT detection for mediating attractive responses. Testing these, as well as many other possibilities, to link semiochemicals to behavioral and physiological outputs, at the level of molecules, cells, and circuits, remain outstanding.

## Methods

### Animal husbandry

Wild-type C57BL/6J (Jackson Labs 000664) and *Bax^-/-^* (Jackson Labs 002994) were bought and maintained at institutional facilities. Procedures of animal handling and tissue harvesting were approved by the Institutional Animal Care and Use Committee of Duke University. Animals were killed within 7 days of the ages reported in this study. Sex-separated male and female mice were socially housed with 2-5 animals per cage. Sex-combined cages contained 1 male and 1 female. All sex-combined cages produced litters. Pups were aged to P21-P28 before being weaned or used for independent experiments.

3 male and 3 female biological replicates were used in each condition to sequence wild-type and *Bax^-/-^* whole olfactory mucosa tissues (MOE + other tissues in the nose). 3 male and 3 female wild-type mice were used in each condition to examine MOE *in situ*. 2-3 male and female Bax^-/-^ mice were used to examine MOE *in situ*. 2-6 sections per mouse were imaged, quantified, and reported as individual data points for each condition.

### Preparation of olfactory tissues for RNA-Seq

Whole olfactory mucosa was rapidly collected in 5 mL tubes and flash-frozen in liquid nitrogen from mice killed by CO_2_ asphyxiation and decapitation. Tissues were kept frozen at −80°C until time of RNA extraction. To extract RNA, 1 mL of TRIzol (Life Technologies 15596026) was added to frozen tissue followed by homogenization until no large pieces were readily identifiable. Homogenized tissue was transferred to new 1.5 mL tubes and centrifuged at max speed for 10 minutes. Supernatant was then transferred to new 1.5 mL tubes containing 0.2 mL chloroform and vortexed for 3 minutes. Samples were again centrifuged at max speed for 15 minutes and the aqueous phase was transferred to new tubes containing 0.5 mL of isopropanol. Samples were incubated at room temperature for 5 minutes and then centrifuged at max speed for 10 minutes. Supernatant was decanted and the visible pellet was washed 150 μL of 75% ethanol, centrifuged, and washed again with 180 μL of 75% ethanol. After centrifugation, ethanol wash was pipetted away and RNA pellets were allowed to air-dry with tube lids kept open for 10 minutes. Pellets were then dissolved in RNase-free water by heating to 55°C for 10 minutes. RNA concentration was quantified using a QUBIT HS RNA Assay Kit (Q32855).

88 μL of sample was subjected to RNase-free DNaseI treatment by the addition of 10 μL of 10X Buffer and 2 μL of RNase-free DNaseI (Roche 04 716 728 001) for 30 minutes at 37°C. Following DNaseI treatment, samples were subjected to a modified RNeasy mini protocol for RNA cleanup (Qiagen 74104). 350 μL of buffer RLT was added to the 100 μL sample, mixed and centrifuged. Then, 250 μL ethanol was added, mixed, and immediately transferred to a mini-column. Sample loaded columns were centrifuged for 30 seconds. 500 μL of ethanol diluted buffer RPE was then used to wash the column twice, and sample was eluted in new 1.5 mL tubes with 100 μL of RNase-free water. Presence of RNA was confirmed by the QUBIT HS RNA Assay Kit.

Amplified cDNA from RNA was prepared using a SMART-Seq v4 Ultra Low Input RNA Kit (Takara 634898) protocol as per the manufacturer’s guidelines. In the case of whole olfactory epithelium sequencing, 2 rounds of cDNA amplification were used with 1000 ng of input RNA. DNA libraries were prepared using a half-sized Nexterra XT DNA Library Preparation Kit (Illumina 15032354) protocol as per the manufacturer’s guidelines. Libraries were sequenced on either HiSeq 2000/2500 (50 base pair single read mode) or NextSeq 500 (75 base pair single read mode) with 6-12 pooled indexed libraries per lane. Reads were aligned and quantified using STAR and RSEM using custom-written code allowing up to 10 read multi-mapping events per transcript. Differential expression analysis was performed with custom-written code in R using a combination of DESeq and EdgeR. Intact Olfrs were filtered, and p-values were then re-corrected by FDR.

### Cloning of ORs and generation of anti-sense RNA probes

Mouse ORs were cloned with sequence information from NCBI. OR ORFs were amplified from genomic DNA using Phusion (ThermoFisher F530S) as per the manufacturer’s guidelines. Amplified fragments were cloned into pCI expression vectors (Promega E1731) containing the first 20 residues of human rhodopsin (Rho-pCI) and were verified by sequencing.

To generate anti-sense digoxigenin (DIG)-RNA probes, ORFs were amplified (Qiagen 203203) from Rho-pCI vectors and purified via a MinElute Kit (Qiagen 28004) using manufacturer protocols with an added T3 polymerase promoter sequence at the 3’ end. Anti-sense RNA was then *in vitro* transcribed using a T3 RNA polymerase (Promega P2083) and a DIG RNA labeling mix (Roche 11277073910) using manufacturer protocols. RNA probes were then alkaline hydrolyzed (80mM NaHCO_3_, 120mM Na_2_CO_3_) for 60°C for 15 minutes and purified using a microcolumn (Bio-Rad 732-6223). Probe integrity was assessed by agarose gel and kept at −80°C when not in use.

To determine the specificity of OR-specific mRNA probes, coding sequences of ORs were retrieved from NCBI Nucleotide and compared to other transcripts in the mouse by NCBI BLAST using the RefSeq RNA database. Only Olfr983 exhibited relatively high similarity to other ORs by this method. OSNs expressing *Olfr983* were therefore determined by visual identification of the brightest and most intense cells positive for the anti-sense *Olfr983* mRNA. Data shown probing for *Olfr910/912* by *in situ* mRNA hybridization was done by anti-sense probes generated against *Olfr910*. Early experiments also using probes against *Olfr912* indicated similar results (not shown).

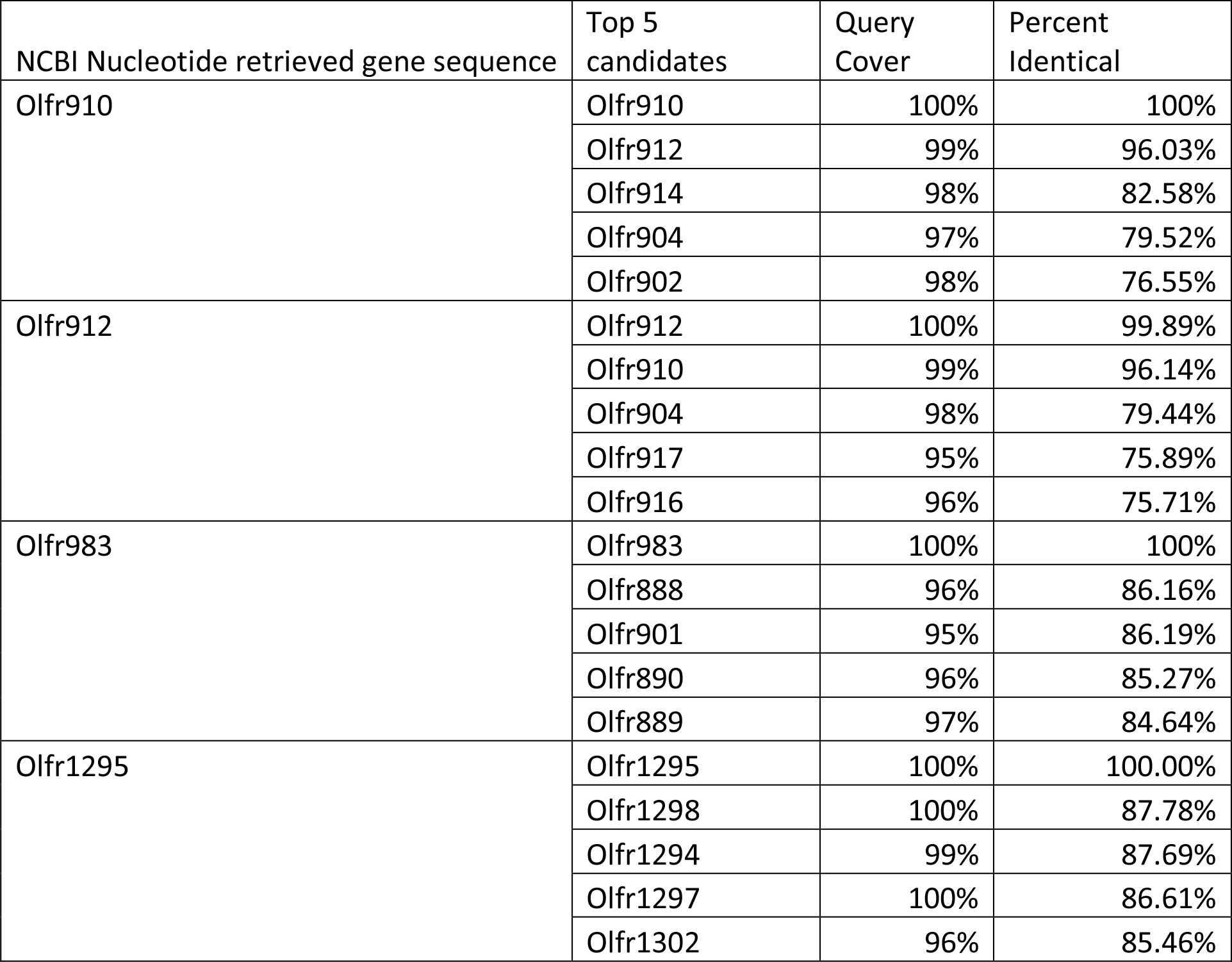

### Preparation of olfactory tissues for staining and *in situ* hybridization

Olfactory epithelium was rapidly dissected and frozen in embedding medium (Tissue-Tek O.C.T. Compound 4583) from mice killed by CO_2_ asphyxiation and decapitation. 18-22 μm fresh frozen coronal sections were cut using a cryostat (Leica CM1850) onto microscope slides (Fisherbrand Superfrost Plus 1255015) and kept at −80°C until use.

For *in situ* RNA probe hybridization, slides were brought to room temperature, dried and rapidly fixed in 4% paraformaldehyde in 1x PBS (pH ∼7.5) for 15 minutes. Slides were then washed twice 1x PBS and submerged into a triethanolamine solution consisting of 700 mL dH_2_O with 8.2 mL triethanolamine. 1.75 mL of acetic anhydride was then added dropwise over the course of 7 minutes with constant and slow stirring for a total of 10 minutes, all at room temperature. Slides were then washed with 1x PBS and blocked with prehybridization solution (see components below) for 1 hour at 58°C in a humidified hybridization oven. RNA probe concentrations were then individually optimized by dilution in prehybridization buffer and pipetted directly onto slides and covered with laboratory film (Parafilm 54956) for overnight hybridization at 58°C. Slides were then rinsed the next day in 72°C heated 5x SSC twice, washed twice in 72°C heated 0.2x SSC for 30 minutes per wash, and again finally washed in 1x PBS for a minimum of 5 minutes at room temperature. Slides were then blocked with 0.5% nucleic acid blocking reagent (Roche 11096176001) dissolved in a 1x PBS containing maleic acid (Sigma M0375) for 30 minutes. Blocking solution was then replaced with 1:1000 horseradish peroxidase (HRP)-conjugated anti-DIG antibody (Roche 11207733910) solution diluted in the blocking medium for 45 minutes. Slides were then washed in 1x PBS three times, 10 minutes per wash, and coated with 0.1% BSA in 1x PBS. Hybridization signals were detected using tyramide signal amplification (TSA) using fluorescein (PerkinElmer) as the fluorophore diluted in 1x PBS containing 0.003% H_2_O_2_ via incubation for 10 minutes in darkness, all at room temperature.

For pS6 immunostaining, slides were blocked in 5% skim milk dissolved in 1x PBS containing 0.1% Triton X-100 at room temperature for 1 hour. Blocking solution was then replaced with 1:300 anti-pS6 antibody (ThermoFisher 44-923G) dissolved in blocking solution and incubated overnight at 4°C. Anti-pS6 antibody was detected using a 1:200 Cy3-conjugated secondary (Jackson Immuno 711-165-152) diluted in 5% skim milk dissolved in 1x PBS by incubation for 45 minutes in darkness. Cell nuclei were detected using a 1/10000 dilution of a 1% bisbenzimide (Sigma bisbenzimide H 33258) solution by incubation for 5 minutes at room temperature. All slides were rinsed in dH2O, cover slipped, and allowed to dry before examination under a microscope.

### Slide microscopy

Z-stacked images with 2-µm intervals between each slice were obtained at 200× magnification using the Zeiss Axiocam MRm and upright inverted fluorescent microscope with ApoTome functionality. The filter sets used were as follows: Zeiss filter set #38 for fluorescein, #43 for Cy3, and #49 for bisbenzimide. For pS6 signal intensity quantification, Cy3 signals (pS6 intensity) within fluorescein positive cells (OR expression) were merged as a maximum intensity projection in ImageJ. Average pS6 signal intensities within single cells were then normalized by average pS6 signal intensities across all OSNs within the same image to report a corrected pS6 signal intensity.

### Odor exposures

All juvenile mice used in this study were approximately 3 weeks old. Mice were habituated for 1 hour in a clean and covered single-use paper container (International Paper DFM85) in a fume hood. For odor exposure, mice were then transferred to a new paper container containing either the mature male mice (∼ 8-weeks-old), mature female mice (∼8-weeks-old), or diluted odorant for another hour. All odorant exposures in this study consisted of 10 μL of stimulus spotted onto a cut blotting pad (VWR 28298-014) placed inside an odor cassette (Tissue-Tek 0006772-01). All odorants unless otherwise stated were diluted in water, vortexed, and rapidly spotted. MTMT was diluted in ethanol. Control experiments for odor exposures consisted of exposing mice to water or ethanol spotted on blotting pads placed in odor cassettes. Odor exposure experiments on juvenile mice employed both male and female mice. Odor exposure experiments on mature mice used male and female mice at or near estrous determined by vaginal cell cytology.

### Phosphorylated S6 ribosomal capture (pS6-IP)

Mice used for pS6-IP were ∼3 weeks old, mixed sex, littermates. Mice were killed by CO_2_ asphyxiation and cervical dislocation. Olfactory tissue was rapidly dissected in Buffer B (2.5mM HEPES KOH pH 7.4, % glucose, 100 μg/mL cycloheximide, 5 mM sodium fluoride, 1 mM sodium orthovanadate, 1 mM sodium pyrophosphate, 1 mM β-glycerophosphate, in Hank’s balanced salt solution). Tissue pieces were then minced in 1.35 mL Buffer C (150 mM KCl, 5 mM MgCl_2_, 10 mM HEPES KOH pH 7.4, 0.100 μM Calyculin A, 2 mM DTT, 100 U/mL RNAsin, 100 mg/mL, 100 μg/mL cycloheximide, protease inhibitor cocktail, 5 mM sodium fluoride, 1 mM sodium orthovanadate, 1 mM sodium pyrophosphate, 1 mM β-glycerophosphate)and subsequently transferred to homogenization tubes for steady homogenization at 250 rpm three times and at 750 rpm nine times at 4°C. Samples were then transferred to a 1.5 mL LoBind tube (Eppendorf 022431021) and clarified at 2000xg for 10 minutes at 4°C. The low-speed supernatant was transferred to a new tube on ice, and to this solution was added 90 μL of NP40 (Sigma 11332473001) and 90 μL of 1,2-diheptanoyl-sn-glycero-3-phosphocholine (DHPC, Avanti Polar Lipids: 100 mg/0.69 ml). This solution was mixed and then clarified at a max speed (17000xg) for 10 minutes at 4C. The resulting high-speed supernatant was transferred to a new tube where 20 μL was saved and transferred to a tube containing 350 μL buffer RLT. To the remainder of the sample, 1.3 μL of 100 mg/mL cycloheximide, 27 μL of phosphatase inhibitor cocktail (250 mM sodium fluoride, 50 mM sodium orthovanadate, 50 mM sodium pyrophosphate, 50 mM β-glycerophosphate) and 6μL of anti-pS6 antibody (Cell Signaling D68F8) was added. The sample was gently rotated for 90 minutes at 4°C. To prepare beads, 100 μL of beads (Invitrogen 10002D) was washed 3 times with 900 μL of buffer A (150 mM KCl, 5 mM MgCl_2_, 10 mM HEPES KOH pH 7.4, 10% NP40, 10% BSA), and once with 500 μL of buffer C. Sample homogenate was added to the beads and incubated with gentle rotation for 60 minutes at 4°C. Following incubation, beads were washed with 4 times with 700 μL of buffer D (350 mM KCl, 5 mM MgCl_2_, 10 mM HEPES KOH pH 7.4, 10% NP40, 2 mM DTT, 100 U/mL RNAsin, 100 μg/mL cycloheximide, 5 mM sodium fluoride, 1 mM sodium orthovanadate, 1 mM sodium pyrophosphate, 1 mM β-glycerophosphate). During the final wash, beads were moved to room temperature, wash buffer was removed, and 350 mL of buffer RLT was added. Beads were incubated in buffer RLT for 5 minutes at room temperature. Buffer RLT containing immunoprecipitated RNA was then eluted and stored at −80°C until clean up using a kit (Qiagen 154025593). cDNA was generated using 11 rounds of amplification with 10 ng RNA input.

### Source of odors

MTMT was synthesized as previously described in Lin et al. 2005^18^. Racemic SBT was synthesized in two steps from 2-aminoethanol and methyl 2-methylbutanedithioate^44^ in >80% yield according to the procedure of Abrunhosa et al. 2001^45^. The final product showed high purity by gas chromatography, ^1^H and ^13^C NMR, and mass spectroscopic data; excellent agreement with reported parameters for this compound in Tashiro et al. 1999^46^. DHB was synthesized as previously described in Wiesler et al. 1984^47^. β-caryophyllene (Sigma W225207), 2,5-DMP (Sigma 175420), 2-heptanone (Sigma W254401), (E)-β-farnesene (Bedoukian P3500-90) were purchased.

## Acknowledgements

We thank Claire A. de March, Ha Na Choe, Conan Juan, and Yen Dinh for comments on the manuscript.

---

**Supplementary Figure 1.**
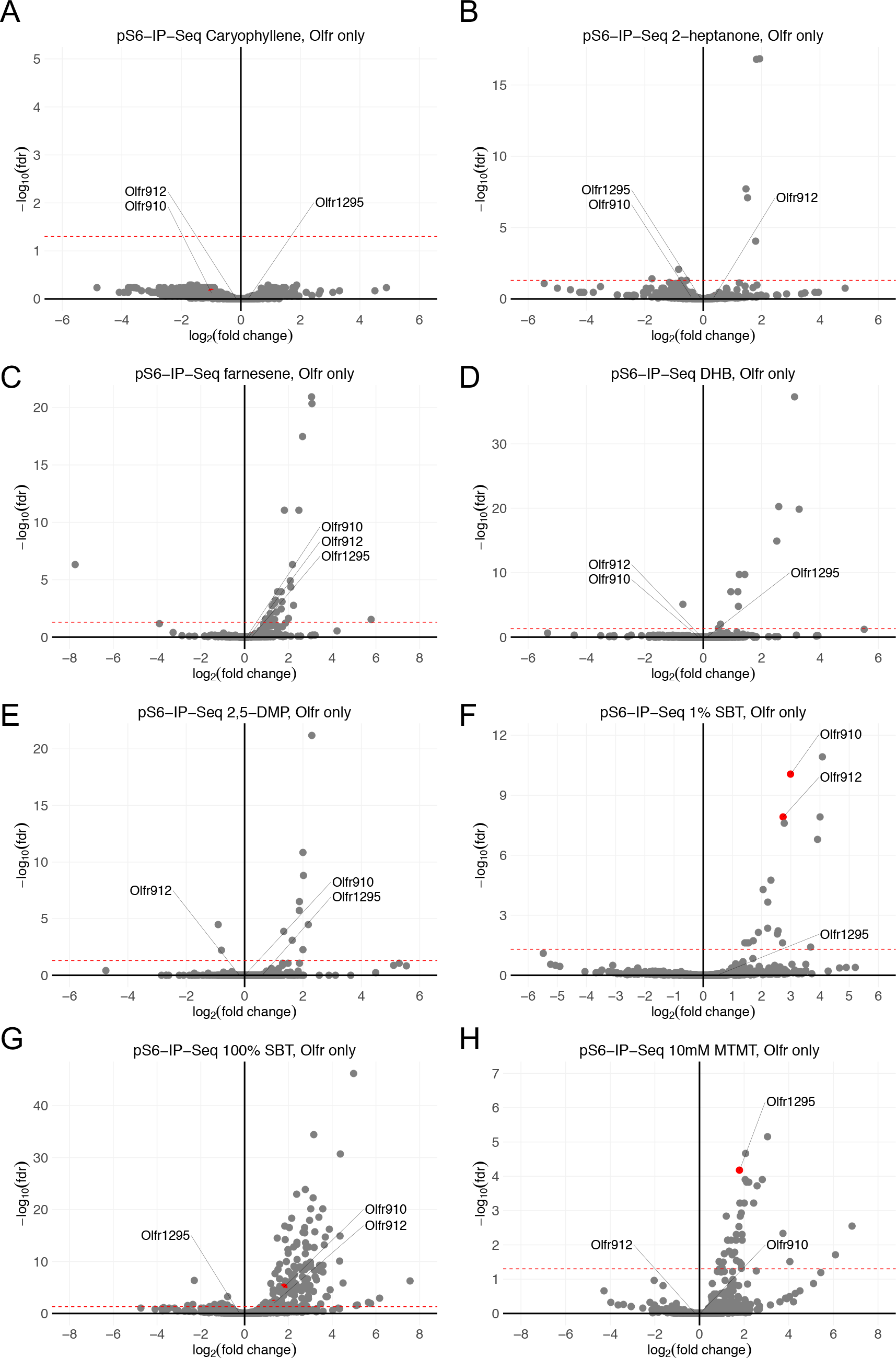
Volcano plots showing the results of pS6-IP-Seq against the battery of odorants. The red dashed line indicates an FDR = 0.05. For SBT and MTMT, pS6-IP-Seq results of concentrations higher than the ones reported in Figure 4 are shown as well. Among the tested odorants, only SBT and MTMT exposure lead to the enrichment of transcripts for *Olfr910*, *Olfr912*, and *Olfr1295*.

**Supplementary Figure 2.**
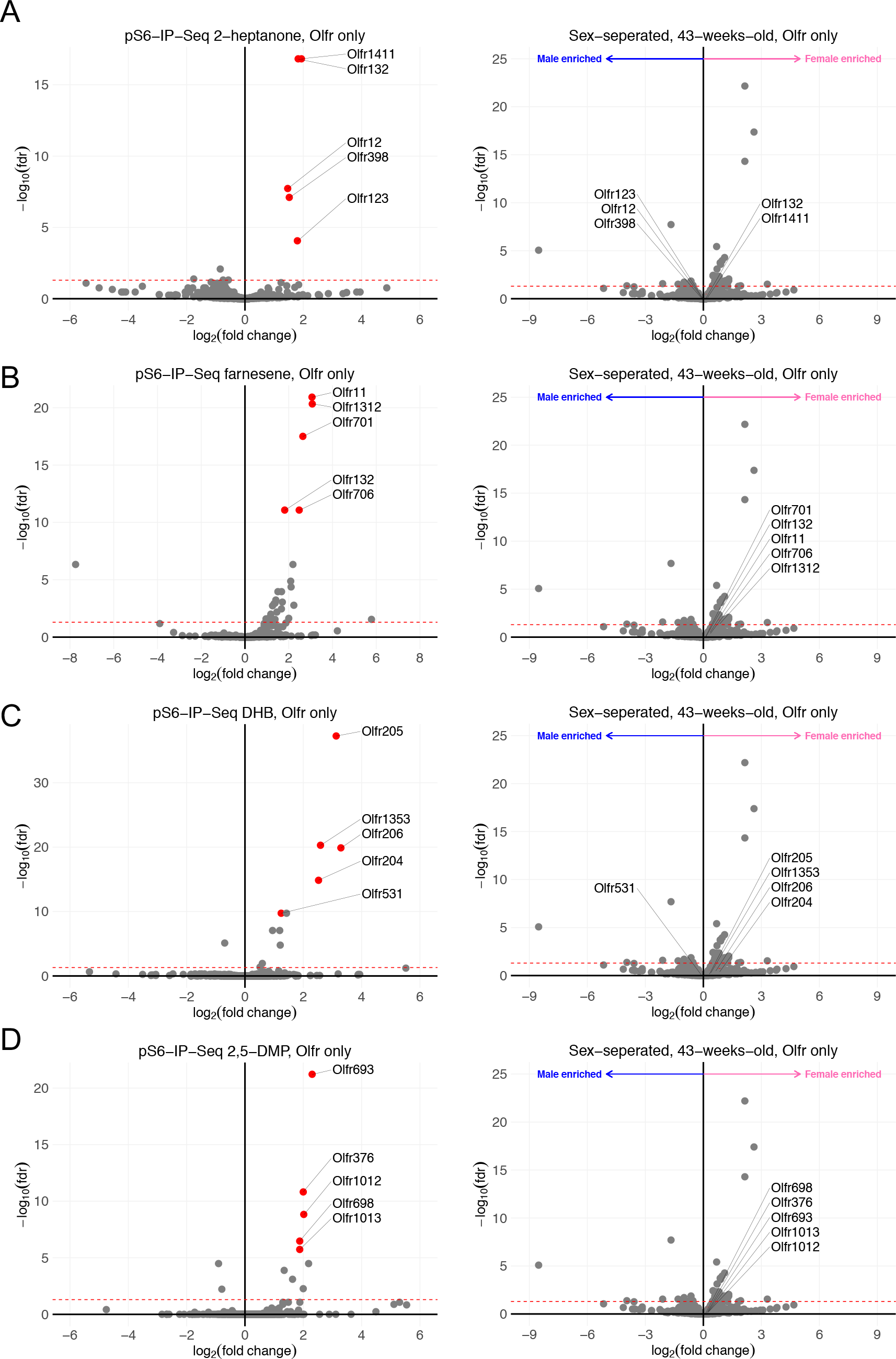
To show that sex-specific receptor expression is limited to ORs robustly responsive to SBT and MTMT, the top 5 candidate receptors for 3,4-dehydro-*exo*-brevicomin, 2,5-dimethylpyrazine, (E)-β-farnesene, and 2-heptanone are highlighted and shown to not exhibit sexually dimorphic expression.

**Supplementary Figure 3.**
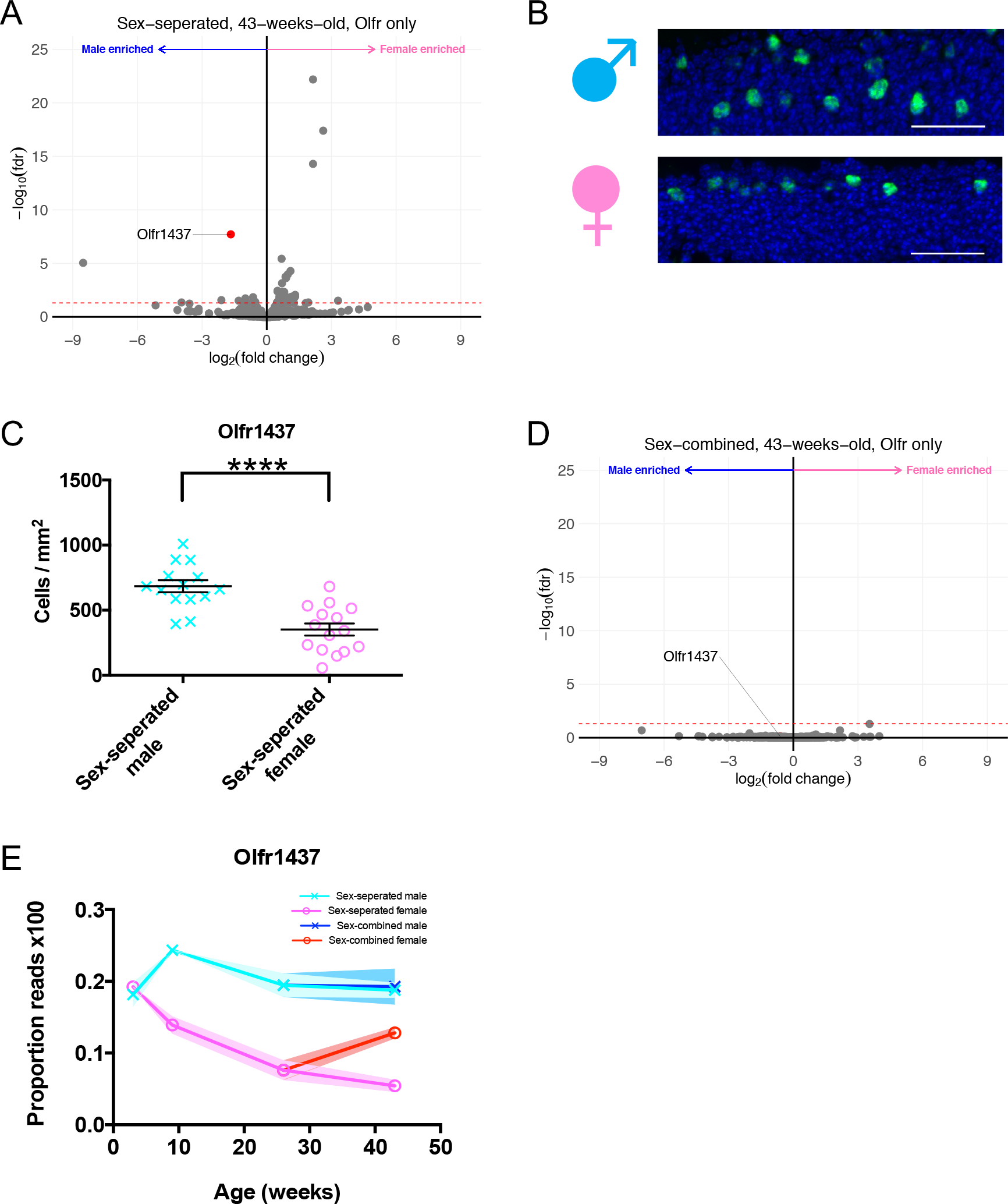
To show our observations of sexual dimorphism in the MOE are not a sex-biased observation, we investigated *Olfr1437*, a male-enriched OR. A. Volcano plot comparing expression of *Olfrs* between 43-week-old sex-separated male and female mice. The red dashed line indicates an FDR = 0.05. *Olfr1437*, a male-enriched OR is highlighted. B. Representative *in situ* hybridization picture probing for the expression of *Olfr1437* in 43-week-old sex-separated male (top) and female (bottom) mice. Scale bars indicate 50 μm. C. Summary data showing the proportion of OSNs expressing *Olfr1437* between 43-week-old sex-separated male and female mice. An unpaired two-tailed t-test revealed statistical difference (p < 0.0001) between males and females. D. Volcano plot comparing expression of *Olfrs* between 43-week-old sex-combined male and female mice. The red dashed line indicates an FDR = 0.05. *Olfr1437*, a male-enriched OR is highlighted. E. Longitudinal plotting of the proportions of reads aligned to *Olfr1437*. Proportions were calculated by comparing reads mapped to *Olfr1437* compared to those mapped to other *Olfrs*.

